# Brain representation in conscious and unconscious vision

**DOI:** 10.1101/2024.05.27.596053

**Authors:** Ning Mei, David Soto

**Author notes:** Correspondence to: Ning Mei; David Soto.

## Abstract

The development of robust frameworks to understand how the human brain represents conscious and unconscious perceptual contents is paramount to make progress in the neuroscience of consciousness. Recent functional MRI studies using multi-voxel pattern classification analyses showed that unconscious contents could be decoded from brain activity patterns. However, decoding does not imply a full understanding of neural representations. Here we re-analysed data from a high-precision fMRI study coupled with representational similarity analysis based on convolutional neural network models to provide a detailed information-based approach to neural representations of both unconscious and conscious perceptual content. The results showed that computer vision model representations strongly predicted brain responses in ventral visual cortex and in fronto-parietal regions to both conscious and unconscious contents. Moreover, this pattern of results generalised when the models were trained and tested with different participants. Remarkably, these observations results held even when the analysis was restricted to observers that showed null perceptual sensitivity. In light of the highly distributed brain representation of unconscious information, we suggest that the functional role of fronto-parietal cortex in conscious perception is unlikely to be related to the broadcasting of information, as proposed by the global neuronal workspace theory, and may instead relate to the generation of meta-representations as proposed by higher-order theories.

## Introduction

The neuroscience of visual consciousness seeks to elucidate the neural underpinnings of subjective experience. To achieve this goal, it is paramount to develop robust frameworks for distinguishing unconscious and conscious information processing. Our study embraces an information-based strategy, integrating machine learning classification approaches and computational modeling techniques with functional Magnetic Resonance Imaging (fMRI) data to understand the properties of unconscious representations in the human brain.

The extent to which conscious and unconscious visual contents are processed in the higher-level stages of the ventral visual stream and associated parieto-prefrontal areas remains the subject of ongoing debate (Boly et al., 2017; Brown et al., 2019; Dehaene, 2014; Lau, 2022). Previous neuroimaging work has shown that object categories of visible stimuli are encoded in the ventral-temporal cortex (Kriegeskorte, 2011; Naselaris et al., 2011) and perceptual and mnemonic contents can also be decoded in fronto-parietal cortex (Christophel et al., 2012; Ester et al., 2015; Kapoor et al., 2020; Panagiotaropoulos et al., 2012).

It has been argued that unconscious contents are temporarily and locally encoded in the visual cortex (Lamme, 2020), while visual consciousness may be associated with activity in large-scale association networks, involving fronto-parietal cortex (cf. the global neuronal workspace model, (Dehaene & Changeux, 2011). However, recent research suggests that unconscious information processing might support higher-order cognition (Berkovitch & Dehaene, 2019; Chong et al., 2014; Güldener et al., 2022; Lau & Passingham, 2007; Rosenthal et al., 2016; Soto et al., 2011; Trübutschek et al., 2017; Van Gaal & Lamme, 2012; Van Gaal et al., 2014; Wuethrich et al., 2018). Nevertheless, the purported influence of unconsciously processed information on behavioural responses has proven challenging to replicate (Stein et al., 2020a). Unconscious processing research has been heavily criticised on methodological grounds related to the low number of trials used in objective awareness tests to exclude conscious awareness, and the use of subjective measures of awareness that are known to be influenced by criterion biases (Newell & Shanks, 2014; Shanks et al., 2021).

One reason for the ongoing controversy around the scope of unconscious information processing is the absence of robust frameworks to effectively isolate the influence of unconscious contents at behavioural and neural levels (Soto et al., 2019; Stein et al., 2020b). Research on unconscious processing is heavily constrained by the low signal to noise ratio of the unconscious stimulus, which is typically presented very briefly and/or strongly masked. If the unconscious item is poorly represented in the brain, it is likely that unconscious effects on behaviour are only weak or difficult to detect (Lau, 2022). However, the absence of observable effects on behavior does not refute the existence of unconscious processing. The brain-based framework to unconscious processing (Soto et al., 2019) suggests that information-based analysis of brain activity patterns can provide insights into the representations of unconsciously processed stimuli, even with null effects on behavioral measures.

A recent fMRI study employing a high-precision, within-participant approach to investigate the neural representation of unconscious contents (Mei et al., 2022b) showed that unconscious contents, even those associated with null sensitivity, can be reliably decoded from multi-voxel patterns that are highly distributed along the ventral visual pathway and also involving parieto-frontal substrates. It should be noted, however, that the capacity to decode specific information does not equate to a comprehensive understanding of the neural code (Kriegeskorte & Douglas, 2018). Hence, beyond linear decodability, pattern classification approaches fall short to delineate the specific representations encoded in brain regions.

Here we re-analysed fMRI data from a previous study (Mei et al., 2022b) in which participants (N = 7) underwent six fMRI sessions across six days while performing a discrimination task for animate and inanimate images presented very briefly and masked. In addition to multi-voxel pattern classification, we used model-based representational similarity analysis (RSA) to provide a fine-grained information-based approach to the neural representation of unconscious and conscious contents. RSA can be used to quantify the geometry of multi-voxel patterns in a given brain region relative to a computational model (Diedrichsen & Kriegeskorte, 2017; Kriegeskorte et al., 2008). In order to model the brain responses to the visual images, both seen and unseen, we assessed the correspondence of the multi-voxel response patterns elicited by the images and the representation of the images given by convolutional neural network models (CNNs), which have been successfully used to explain brain activity in object recognition tasks (Güçlü & van Gerven, 2015; Khaligh-Razavi & Kriegeskorte, 2014; Kriegeskorte, 2009; Schrimpf et al., 2020; Seeliger et al., 2018).

## Methods

### Experimental paradigm

Participants were asked to discriminate whether the target images were animate or inanimate (Moreno-Martínez & Montoro, 2012). The images images were presented briefly, preceded and followed by a dynamic mask made of Gaussian noise. On each trial of the fMRI experiment, participants were required to discriminate the image category and to rate their awareness using the following response categories: (i) no experience/just guessing (ii) brief glimpse (iii) clear experience with a confident response. The duration of the images was based on an adaptive staircase that was running throughout the experiment, and which, based on pilot testing, was devised to obtain a high proportion of unconscious trials. On average, target duration was (i) 25 ms on trials rated as unaware (ii) 38 ms on glimpse trials and (iii) 47 ms on the aware trials. Figure 1 illustrates an example of a trial in the behavioural task. Additional details can be found in Mei et al. (2022). (Mei et al., 2022b)

**Figure 1:**
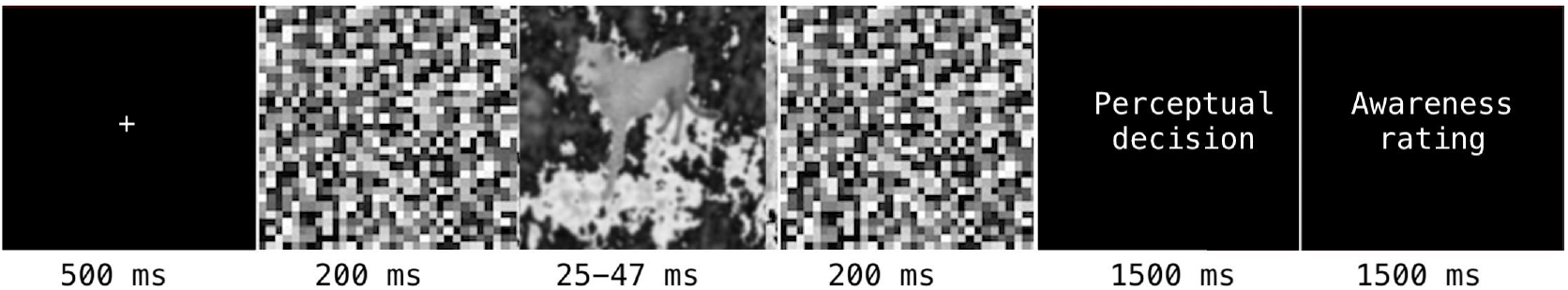
Experimental paradigm. Example of the sequence of events within an experimental trial. Participants were asked to discriminate the category of the masked image (animate v.s. inanimate) and then rate their visual awareness on a trial by trial basis.

### Encoding-based RSA pipeline

Standard RSA characterizes the structure of cognitive or perceptual representations within a feature space in terms of distances between response vectors associated with different aspects of the task or the experimental stimuli (Kriegeskorte et al., 2008). This is measured by distance-based measurements such as Euclidean distance (Kriegeskorte et al., 2006) between neural activity patterns for each pair of experimental conditions or for each pair of representations given by a computational model. Konkle and colleagues (Konkle & Alvarez, 2022) proposed an encoding-based RSA pipeline in which encoding models are added on top of the standard RSA pipeline. The encoding models help to better contextualize how the representations given by computational models (i.e. convolutional neural networks, such as ResNet50 (He et al., 2016)) relate to the brain representations. In order to allow for robust statistical inference in our highly-sampled within-subject design, the encoding model of the RSA pipeline was fitted and tested across pairs of participants. Standard RSA pipelines are not designed to perform within-subject statistical inference. Descriptive results from exploratory RSA pipelines assessed in a within-subject manner are reported in Mei et al. (2022a). Here we present an improved encoding-based RSA framework using a cross-participant cross-validation approach. This approach produces independent RSA maps across participants that could be fed to statistical permutation tests.

#### Extraction of the representations of the images from computer vision models

ResNet50 was specifically chosen for RSA given its superior performance among the evaluated computer vision models in the Brain-Score challenge (Schrimpf et al., 2018; Schrimpf et al., 2020). The “50” in ResNet50 denotes the approximate depth of the network (i.e. around 50 layers). ResNet50’s architecture is characterized by its repetitive use of residual blocks with skip connections, enabling the training of very deep networks without degradation problems, and achieving high performance in image recognition tasks.

First, we fine-tuned the pre-trained ResNet50 architecture (He et al., 2016), which was initially trained on ImageNet (Deng et al., 2009), with the Caltech101 dataset (Fei-Fei et al., 2004) to reduce the dimensionality of hidden representations to 300. During fine-tuning, the pretrained ResNet50 (He et al., 2016) was stripped of the original fully-connected layer while weights and biases of the convolutional layers were frozen and not updated further (Yosinski et al., 2014). An adaptive pooling (McFee et al., 2018) operation was applied to the last convolutional layer so that its output became a onedimensional vector, and a new fully-connected layer took the weighted sum of the previous outputs (i.e. the ‘hidden layer’). The outputs of the hidden layer were passed to a Scaled Exponential Linear Unit (SELU) activation function (Klambauer et al., 2017).

A new fully-connected layer, namely, the classification layer, took the outputs processed by the activation function of the hidden layer to compose the classification layer. The model was trained using 96 unique categories of the Caltech101 images (BACKGROUND_Google, Faces, Faces_easy, stop_sign, and yin_yang were excluded). The convolutional layers were frozen during the training, while the weights of the newly added two layers were modified. The loss function was binary cross entropy and the optimizer was Stochastic Gradient Descent. The data was split into train and validation partitions, and the training was terminated if the performance on the validation data did not improve for five consecutive epochs.

We then fed the trained FCNNs with exactly the same images used in our experiment but without the noise background, in order to extract the hidden representations of the images. We then averaged the hidden representations of the trials belonging to the same item (i.e cat) and computed the model RDMs for the ResNet50 (Figures 2 depict the RDM for ResNet50 hidden representations), where they appear clear clusters for the animate and inanimate image categories. The similarities were more consistent among animate images (lower left quadrant) compared to the inanimate images (upper right quadrant).

**Figure 2:**
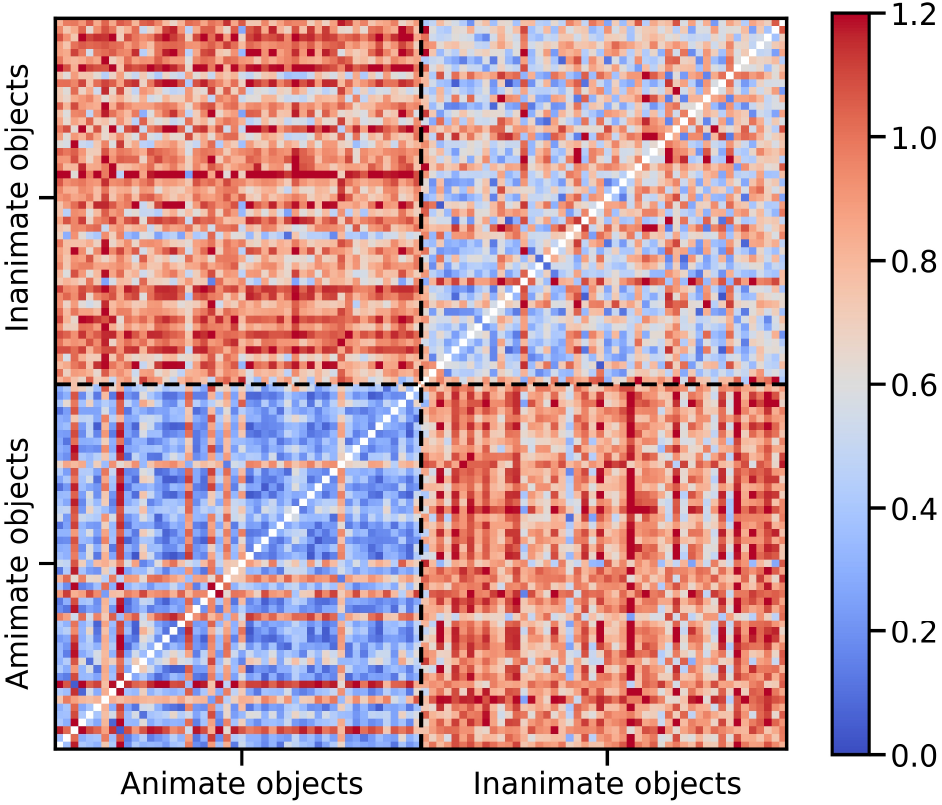
Representational dissimilarity matrix of the hidden representations of the ResNet50 model fine-tuned by Caltech101 dataset. The images were resized to 128 × 128 × 3 and then passed through the model to obtain the feature representations. The RDM was computed by 1 - Pearson correlations of the feature representations. The first 48 items were animate and the last 48 items were inanimate.

### Cross-validation of the encoding-based RSA across participants

A standard RSA pipeline involves computing a model representational dissimilary matrix (RDM) and a corresponding brain RDM, and then compute the similarity (i.e. correlation) between the model and the brain RDMs (Kriegeskorte, 2015; Kriegeskorte & Douglas, 2019; Kriegeskorte et al., 2008). The encoding-based RSA (Konkle & Alvarez, 2022) begins by fitting an encoding model to predict brain responses (Konkle & Alvarez, 2022). The RSA is then computed between the predicted brain responses and the true brain responses.

In the present study, the encoding-based whole-brain searchlight RSA was cross-validated across participants. For a given cross-validation fold, trials of a given awareness condition (i.e., conscious) that belonged to a given participant were used to fit a encoding model. The encoding model used CNN features to predict the voxelwise BOLD responses. Subsequently, we applied the encoding model to trials of a given awareness condition (i.e., unconscious) that belonged to another participant. The observed and predicted BOLD responses were averaged over unique items. Thus, the average of the observed and the predicted BOLD responses across unique stimuli formed two 96 × 96 RDMs, and their lower triangular sections were correlated using Spearman correlation for subsequent analysis. The analysis pipeline was executed on the standard MNI space using a searchlight sphere with radius of 6 *mm*. The Spearman correlation coefficients were assigned to the center of the sphere.

Three distinct cross-participant validation scenarios for the encoding-based RSA were employed: (1) within-conscious-state cross-validation, (2) within-unconscious-state cross-validation, and (3) conscious-to-unconscious generalization.

The cross-participant whole-brain searchlight encoding-based RSA produced 42 distinct searchlight maps resulting from the combination of train and test participant pairs. We then assessed the statistical significance of the correlation maps by means of a non-parametric t test against zero using the Randomise tool from FSL (Jenkinson et al., 2012) with Threshold-Free Cluster Enhancement (TFCE) (Smith & Nichols, 2009).

### Decoding pipeline

We performed a searchlight analysis to pinpoint the brain regions whose activity patterns discriminated the animate versus inanimate dimension of the image. We used a similar between-participant cross-validation method as described in the encoding-based RSA methods. Each participant’s fMRI dataset was normalized to the standard MNI space prior to conducting the analysis (Collins et al., 1994).

For a given cross-validation fold, trials of a given awareness condition (i.e., conscious), from a given participant, were used to fit a decoding model. Subsequently, we applied the decoding model to trials of a given awareness condition (i.e., unconscious), from a different participant. The decoding model comprised a scalar for feature normalization and an L1-regularized linear support vector machine (SVM) for classification. During classifier training, the scalar derived means and standard deviations were used to standardize the features. These parameters were locked during the test. We used L1-regularization to facilitate feature selection by assigning zero weights to sparsity—irrelevant features (Vidaurre et al., 2013). Probabilistic predictions made by the SVM on the normalized test data were compared against the actual labels using a Receiver Operating Characteristic Curve (ROC AUC) approach. This cross-validation procedure was run within a moving searchlight sphere with radius of 6*mm*, and the ROC AUC scores were assigned to the center of the spheres.

The decoding analysis also included cross-validation procedures across the different awareness states (i) within the conscious trials, (ii) within the unconscious trials, and (iii) from conscious to unconscious trials. For each cross-validation procedure, 42 brain maps were produced resulting from the combination of subject pairs used for training and testing the decoder. A permutation t-test using Randomise from FSL with TFCE was used to assess the statistical significance of the voxels in which the decoding scores were greater than 0.5.

## Results

The signal detection theoretic analyses of the behavioural data can be found in Mei et al. (2022b). In short, four of the participants showed null perceptual sensitivity of the stimulus category on the trials rated as unaware, while the remaining three participants showed ‘blindsight’ like behaviour (i.e. above chance discrimination performance on trials reported as unaware). In the following presentation of the results we use the label ‘unconscious’ to refer to the trials in which the participants reported to have no experience of the visual stimulus and ‘conscious’ to refer to the trials in which the subjects reported clear experience.

### Encoding-based RSA

Building on the work of Konkle and Alvarez (2022), we incorporated a cross-participant validation approach within an encoding-model-driven RSA framework. We trained the encoding model with the trials of a given awareness condition (i.e., conscious) from a given participant. Then, the model was used to predict the BOLD signals of a different participant in a given awareness condition (i.e., unconscious trials). The predicted BOLD signals, in conjunction with the actual BOLD responses from the test participant, served as inputs for conducting a whole-brain searchlight RSA analysis (see Methods).

In the conscious trials, there was a highly distributed set of regions involving the ventral visual pathway and fronto-parietal areas in which BOLD responses significantly correlated with the computer vision model representations (Figure 3). The generalization from conscious to unconscious trials showed a similar distributed profile (Figure 4), which, remarkably, was also confirmed by the analysis restricted to the unconscious trials (Figure 5).

**Figure 3:**
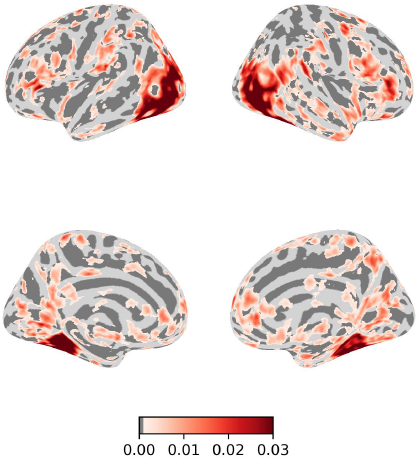
Searchlight RSA results in the conscious trials. The encoding-based RSA was trained on the conscious trials of one participant and tested on the conscious trials of another participant. Spearman correlation coefficients were assigned to the center of the moving searchlight sphere. Clusters are whole-brain corrected (*α* = 0.05)

**Figure 4:**
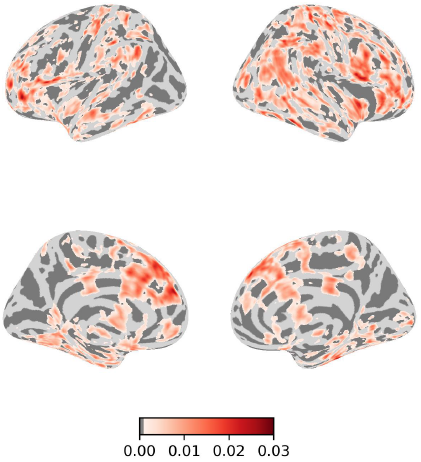
Results from the encoding-based searchlight RSA trained on the conscious trials and tested on the unconscious trials. Spearman correlation coefficients were assigned to the center of the moving searchlight sphere. Clusters are whole-brain corrected (*α* = 0.05)

**Figure 5:**
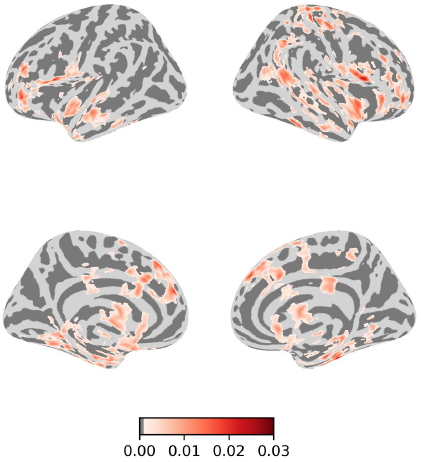
Searchlight RSA results in the unconscious trials. Spearman correlation coefficients were assigned to the center of the moving searchlight sphere. Clusters are whole-brain corrected (*α* = 0.05)

### Decoding results

The searchlight decoding analysis showed that the stimulus category could be decoded from activity patterns in the ventral visual pathway, parietal, and prefrontal cortex. All these areas contained activity patterns that predicted the stimulus category and that generalised awareness states and across participants (Figure 6). The generalization from conscious to unconscious trials showed a similar distributed pattern of brain representation, including the ventral visual pathway, parietal, and prefrontal cortex (Figure 7), which also held when the analysis was performed on the unconscious trials (Figure 8).

**Figure 6:**
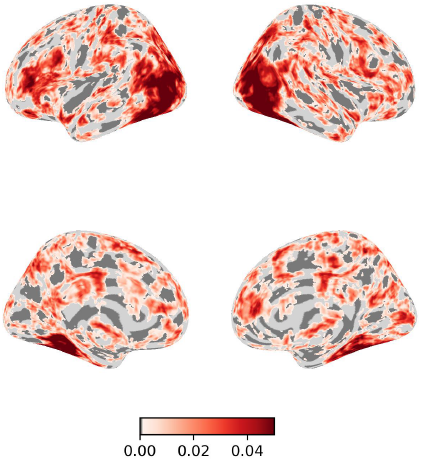
Decoding results within the conscious trials. The classifier was trained using the conscious trials of one participant and tested with the conscious trials of another participant. The decoding scores are shown relative to 0.5. Clusters are whole-brain corrected (*α* = 0.05)

**Figure 7:**
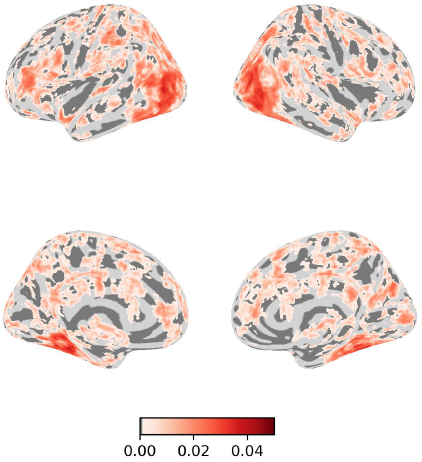
Decoding results. Generalization from conscious to unconscious trials. The classifier was trained using the conscious trials of one participant and tested with the unconscious trials of another participant. The decoding scores are shown relative to 0.5. Clusters are whole-brain corrected (*α* = 0.05)

**Figure 8:**
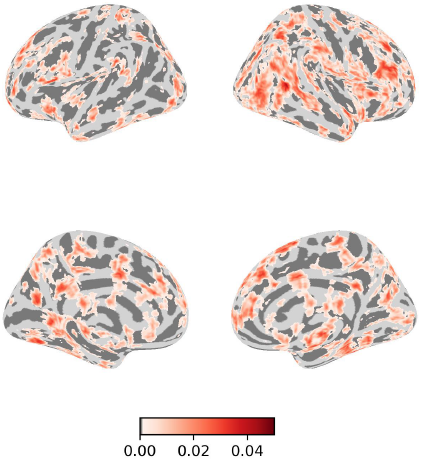
Decoding results within the unconscious trials. The classifier was trained using the unconscious trials of one participant and tested with the unconscious trials of another participant. The decoding scores are shown relative to 0.5. Clusters are whole-brain corrected (*α* = 0.05)

### Encoding-based RSA and decoding analysis for participants who showed null perceptual sensitivity

We then performed the analyses on the four participants that showed null perceptual sensitivity (i.e. no significantly greater than zero in stratified permutation tests; see Mei et al. (2022b)). Accordingly, we excluded the three participants showing above chance perceptual sensitivity on trials in which they reported to be unaware of the target stimulus. In this way, we could rule out the potential influence of the criterion biases in reporting the absence of awareness and hence the possibility that above chance perceptual sensitivity may reflect some degree of “conscious” influence that could also conflate the observed brain activity patterns.

The results are presented in (Figure 9) and (Figure 10). Overall, the pattern of results is similar to the previous analysis with the full sample of participants. This includes distributed activity patterns in ventral temporal and parietofrontal areas. Some of the decoding cross-validation results only showed significant clusters in visual and parietal areas areas. However it should be noted that by reducing the sample size to four the number of folds for the cross-validation was also smaller, thereby reducing the chances of detecting significant results.

**Figure 9:**
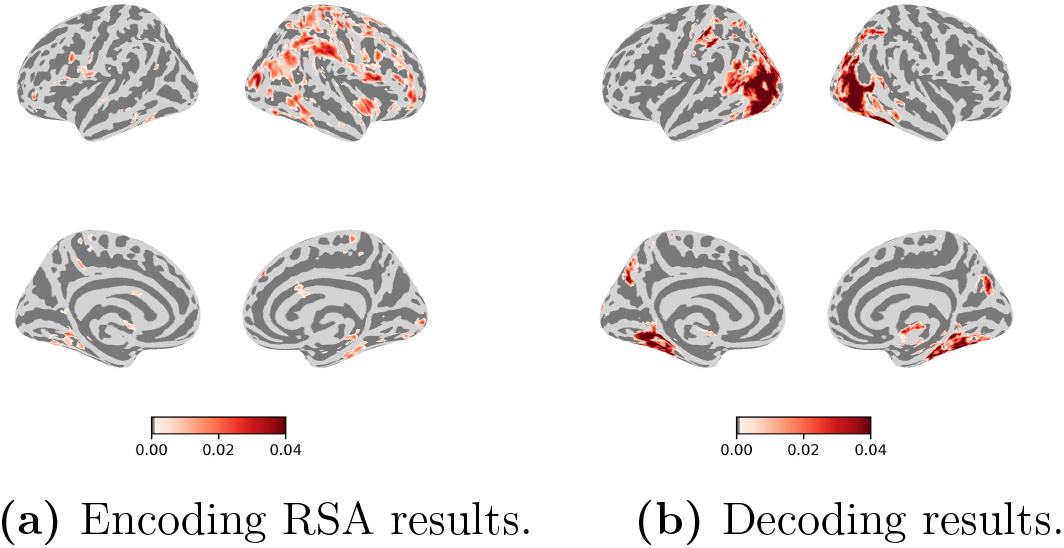
Results from the encoding-based searchlight RSA (a) and decoding (b) on the observers that showed null perceptual sensitivity. Models were trained on the conscious trials and tested on the unconscious trials. Spearman correlation coefficients were assigned to the center of the moving searchlight sphere. Clusters are whole-brain corrected (*α* = 0.05)

**Figure 10:**
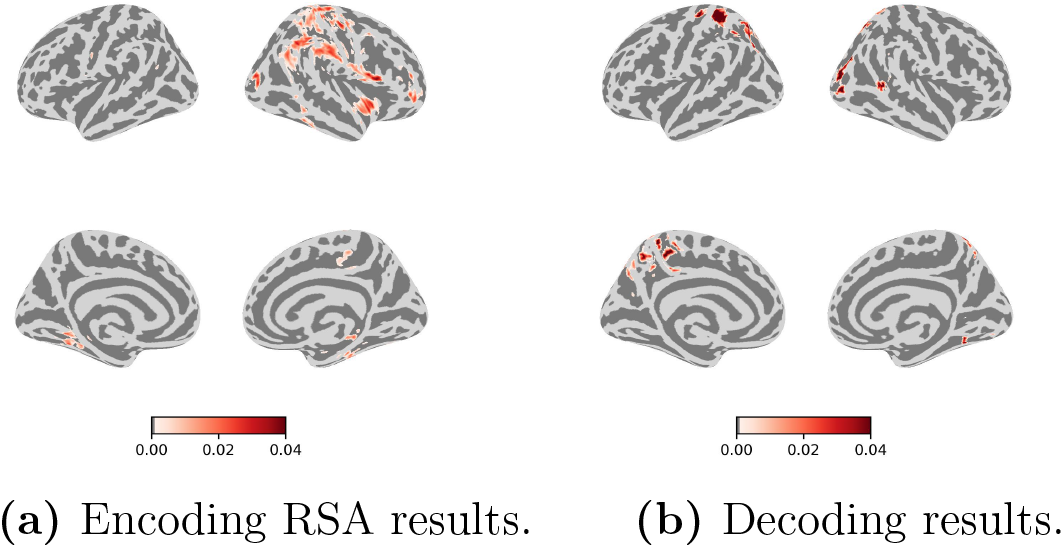
Results from the encoding-based searchlight RSA (a) and decoding (b) in the unconscious trials on the observers that showed null perceptual sensitivity. Spearman correlation coefficients were assigned to the center of the moving searchlight sphere. Clusters are whole-brain corrected (*α* = 0.05)

## Discussion

Developing robust paradigms for assessing the brain representation of unconscious and conscious contents is critical to make progress in the neuroscience of consciousness at both theoretical and empirical levels. Here, we employed an information-based approach, using fMRI in conjunction with model-based RSA and decoding pipelines, to investigate the brain representation of visual objects across different awareness states.

The encoding-based RSA searchlight showed that computer vision model representations strongly correlated with neural responses during object recognition in ventral visual cortex. This result aligns with prior work demonstrating that CNNs are a good computational model of the ventral visual pathway (Güçlü & van Gerven, 2015; Khaligh-Razavi & Kriegeskorte, 2014; Kriegeskorte, 2015; Yamins et al., 2013; Yamins & DiCarlo, 2016). The encoding-based RSA results revealed a highly distributed set of brain regions, including fronto-parietal cortex, involved in the representation of both conscious and unconscious perceptual input. It may be argued that a representational account of brain function also requires that the encoded information guides behavioral performance (Kriegeskorte & Diedrichsen, 2019; Shea, 2018). However, whether or not strongly masked (unconscious) information influences behavior may depend of different factors (e.g. related to brain state) that remain to be determined (Soto et al., 2019). Our pattern of results indicate that unconscious perceptual contents are indeed represented in brain activity, which may have behavioural consequences under certain experimental conditions.

In keeping with the encoding-based RSA, the decoding results showed that, in both conscious and unconscious trials, the stimulus category could be decoded in a highly distributed set of brain regions involving the ventral visual pathway and fronto-parietal substrates. Moreover, the decoding analysis confirmed the significant generalization of a model trained on the conscious trials and tested on the unconscious trials that was also observed in the RSA. This observation is in line with our previous fMRI study using a within-subject analytical approach based on regions of interest (Mei et al., 2022b), in which we showed that unconscious contents are decodable in fusiform, parietal and prefrontal cortex in most of the participants, with significant generalization from conscious to unconscious trials.

These observations are in contrast to prior reports that neural markers of unconscious information processing are restricted to visual cortical regions (Dehaene et al., 2001; Jiang et al., 2007; Ludwig & Hesselmann, 2015; Moutoussis & Zeki, 2002; Stein et al., 2021). However, a few fMRI studies have previously reported prefrontal involvement during unconscious processing in syntactic and semantic tasks (Axelrod et al., 2014; Sheikh et al., 2019), visual short-term memory (Bergström & Eriksson, 2017; Dutta et al., 2014) and cognitive control (Lau & Passingham, 2007; Van Gaal et al., 2008). The increased number of trials per participant in the study by Mei et al. (2022) may well explain the higher sensitivity to decode the content of unconsciously processed information across the brain.

Notably, the decoding and the encoding-based RSA was fitted using brain responses from a given participant and then tested on a different participant. The results from both approaches reveal a notable degree of similarity across participants in the neural representations of conscious and unconscious content. A recent fMRI study using an encoding model with visible images has shown a higher inter-individual variation in the representations in the prefrontal cortex (Lin & Lau, 2024) relative to the ventral visual cortex, which appears more stable across participants. The present results indicate that despite the potential inter-subject variability in the prefrontal representation of visual content, there appears to be a common blueprint in the brain that supports a common, shared representations across participants and states of visual awareness.

The involvement of fronto-parietal cortex in the representation of unconscious content, along with the significant generalization between conscious and unconscious representations, suggests that the functional role of fronto-parietal cortex in conscious perception is unlikely related to the broadcasting of information, as proposed by the global neuronal workspace model (Dehaene et al., 1998). Higher-order theories of consciousness also suggest that the prefrontal cortex is critical for consciousness. However, according to higher-order theories, the role of prefrontal cortex in consciousness relates to the re-representation of the content being represented in downstream sensory regions (i.e. meta-representation). Higher-order theories allow for a significant scope of unconscious processing in brain and behavioural responses, and they do not simply map consciousness with stronger prefrontal signals (Brown et al., 2019). The meta-representational role of the prefrontal cortex that is necessary for consciousness according to the higher-order view likely depends on long-range feedback connections (Huang et al., 2020) and also on other complex neural coding schemes and operations that are difficult to measure with fMRI. Additional work is needed to make further determinations.

## Author contributions

N.M. and D.S. designed the study; N.M. analysed the data; N.M. and D.S. wrote the paper.

## Data availability statement

Analysis scripts are available at https://github.com/nmningmei/unconfeats. The fMRI data can be found at https://openneuro.org/datasets/ds003927

## Acknowledgements

D.S. acknowledges support from the Basque Government through the BERC 2018-2021 program, from the Spanish Ministry of Economy and Competitiveness, through the ‘Severo Ochoa’ Programme for Centres/Units of Excellence in R & D (CEX2020-001010-S) and also from project grants PSI2016-76443-P and PID2019-105494GB-I00 from MINECO.

